## Supplementary Online Material for "Fine-tuning protein language models boosts predictions across diverse tasks"

#### for

### 1 Detailed results - Top-level comparison

Here we provide the results of all individual training runs for using (frozen) pre-trained embeddings (Table S1) and fine-tuning (Table S2). These results are also available in their aggregated form with 95% confidence intervals in Tables S3-S6, which were used to create Fig. 1.

#### 1.1 Individual predictor results

**Table S1.** Individual training runs - pre-trained embeddings

| Model | Rand. Seed | GFP | AAV | GB1 | Stab. | Melt. | Sub. Loc. | Dis. | Sec. Str. |
|---|---|---|---|---|---|---|---|---|---|
| ESM2 8M[1] | 99 | 63.5% | 68.2% | 82.6% | 71.6% | 56.1% | 52.0% | 70.0% | 75.3% |
| | 98 | 63.9% | 66.5% | 81.4% | 75.6% | 57.1% | 53.1% | 70.1% | 75.2% |
| | 97 | 63.8% | 66.3% | 81.7% | 75.1% | 56.8% | 51.6% | 69.3% | 75.2% |
| | 96 | 64.1% | 66.4% | 81.9% | 74.8% | 55.9% | 52.2% | 69.7% | 75.3% |
| | 95 | 64.1% | 66.9% | 81.4% | 78.0% | 56.2% | 51.2% | 69.6% | 75.2% |
| ESM2 35M[1] | 99 | 65.1% | 63.8% | 82.3% | 66.9% | 58.4% | 54.3% | 68.7% | 78.2% |
| | 98 | 65.1% | 59.3% | 82.9% | 67.7% | 58.3% | 54.9% | 67.8% | 78.3% |
| | 97 | 64.7% | 56.9% | 82.5% | 70.0% | 59.0% | 56.7% | 69.1% | 78.2% |
| | 96 | 64.9% | 54.4% | 82.6% | 70.3% | 59.3% | 54.3% | 68.9% | 78.3% |
| | 95 | 65.1% | 60.7% | 83.1% | 70.5% | 58.1% | 54.5% | 68.9% | 78.2% |
| ESM2 150M[1] | 99 | 63.9% | 67.4% | 84.0% | 79.3% | 61.2% | 58.8% | 71.6% | 82.1% |
| | 98 | 63.7% | 65.3% | 83.8% | 75.3% | 61.1% | 60.0% | 71.1% | 82.2% |
| | 97 | 63.9% | 62.9% | 84.0% | 78.7% | 62.2% | 61.0% | 71.3% | 82.0% |
| | 96 | 63.9% | 65.4% | 81.4% | 78.1% | 61.1% | 58.8% | 71.0% | 82.2% |
| | 95 | 63.8% | 67.4% | 83.4% | 80.7% | 62.6% | 58.6% | 70.1% | 82.2% |
| ESM2 650M[1] | 99 | 64.9% | 51.4% | 86.1% | 64.9% | 65.4% | 61.6% | 72.5% | 84.8% |
| | 98 | 64.7% | 45.0% | 85.9% | 64.1% | 65.0% | 64.7% | 71.5% | 84.8% |
| | 97 | 64.7% | 47.7% | 85.9% | 66.3% | 66.6% | 61.8% | 72.2% | 84.8% |
| | 96 | 64.8% | 44.1% | 84.8% | 65.1% | 66.0% | 62.2% | 71.9% | 84.7% |
| | 95 | 64.9% | 47.5% | 85.5% | 71.7% | 65.2% | 64.3% | 72.1% | 84.6% |
| ESM2 3B[1] | 99 | 64.9% | 77.6% | 85.6% | 80.2% | 67.7% | 63.9% | 71.3% | 85.6% |
| | 98 | 64.9% | 77.5% | 86.8% | 78.7% | 67.8% | 63.5% | 70.9% | 85.6% |
| | 97 | 65.0% | 77.3% | 87.1% | 78.4% | 67.6% | 65.1% | 71.3% | 85.5% |
| | 96 | 65.2% | 77.3% | 86.8% | 78.4% | 67.8% | 64.3% | 70.8% | 85.6% |
| | 95 | 65.1% | 78.0% | 86.9% | 78.8% | 67.5% | 64.1% | 70.7% | 85.6% |
| ProtT5[2] | 99 | 61.4% | 70.1% | 85.6% | 77.0% | 68.8% | 62.0% | 71.0% | 84.2% |
| | 98 | 61.6% | 71.0% | 85.2% | 76.9% | 69.6% | 63.1% | 71.0% | 84.3% |
| | 97 | 60.5% | 64.1% | 85.7% | 79.3% | 71.1% | 59.0% | 70.9% | 84.4% |
| | 96 | 61.8% | 67.6% | 84.6% | 76.0% | 68.6% | 60.4% | 70.6% | 84.4% |
| | 95 | 61.4% | 69.8% | 84.8% | 79.6% | 69.0% | 62.2% | 71.1% | 84.3% |
| Ankh base[3] | 99 | 65.9% | 62.2% | 85.3% | 74.0% | 56.3% | 60.0% | 70.9% | 84.4% |
| | 98 | 65.9% | 60.4% | 84.5% | 74.7% | 58.6% | 59.8% | 70.8% | 84.3% |
| | 97 | 65.8% | 62.0% | 85.3% | 73.4% | 57.9% | 61.4% | 70.6% | 84.4% |
| | 96 | 65.9% | 64.2% | 84.6% | 80.5% | 58.1% | 59.8% | 71.4% | 84.3% |
| | 95 | 66.3% | 69.9% | 83.8% | 75.4% | 57.9% | 60.4% | 71.0% | 84.4% |
| Ankh large[3] | 99 | 67.1% | 75.4% | 85.4% | 69.4% | 60.8% | 61.4% | 69.9% | 86.0% |
| | 98 | 67.0% | 71.1% | 86.1% | 65.2% | 62.7% | 58.8% | 69.9% | 86.0% |
| | 97 | 66.8% | 72.7% | 85.7% | 66.4% | 62.2% | 59.6% | 70.0% | 86.0% |
| | 96 | 67.1% | 72.9% | 83.9% | 67.4% | 61.7% | 62.4% | 69.4% | 86.0% |
| | 95 | 67.1% | 72.7% | 85.8% | 77.3% | 62.7% | 60.8% | 70.0% | 86.1% |

For fine-tuning (Table S2) we generally did three reruns for each experiment. The only exception was a clear outlier for full fine-tuning ESM2 150M, *AAV*, random seed 98 which we therefore reran (with random seed 96). The 70.1% result was excluded for all further calculations. For ProtT5 unrelated experiments not published here resulted in additional values for three of the datasets (*Sub-cellular Location, Disorder* and *Secondary Structure* prediction), which we included as well.

**Table S2.** Individual training runs - fine-tuning

| Model | Fine-tuning | Rand. Seed | GFP | AAV | GB1 | Stab. | Melt. | Sub. Loc. | Dis. | Sec. Str. |
|---|---|---|---|---|---|---|---|---|---|---|
| ESM2 8M[1] | full model | 99 | 68.6% | 82.5% | 89.1% | 77.3% | 58.1% | 54.5% | 71.0% | 76.2% |
| | | 98 | 68.9% | 84.9% | 88.2% | 74.6% | 58.5% | 57.1% | 71.6% | 76.2% |
| | | 97 | 69.1% | 80.6% | 87.5% | 77.8% | 58.8% | 55.9% | 71.8% | 76.0% |
| ESM2 35M[1] | full model | 99 | 69.2% | 77.6% | 87.4% | 76.6% | 58.2% | 59.0% | 72.7% | 79.1% |
| | | 98 | 68.7% | 81.8% | 88.1% | 77.4% | 59.0% | 56.5% | 74.3% | 79.1% |
| | | 97 | 68.9% | 83.1% | 88.2% | 76.7% | 62.3% | 58.8% | 73.1% | 79.2% |
| ESM2 150M[1] | full model | 99 | 69.3% | 84.8% | 87.8% | 74.0% | 63.8% | 60.6% | 74.8% | 82.7% |
| | | 98 | 68.9% | 70.1% | 87.8% | 74.1% | 62.6% | 61.4% | 72.9% | 82.9% |
| | | 97 | 69.0% | 84.8% | 88.1% | 71.9% | 61.5% | 62.7% | 74.4% | 82.8% |
| | | 96 | - | 84.2% | - | - | - | - | - | - |
| ESM2 650M[1] | full model | 99 | 68.8% | 78.1% | 87.1% | 69.8% | 66.1% | 65.1% | 72.0% | 85.5% |
| | | 98 | 69.2% | 80.7% | 88.6% | 78.5% | 67.8% | 63.5% | 72.4% | 85.5% |
| | | 97 | 68.5% | 76.2% | 88.1% | 74.5% | 65.3% | 62.9% | 71.6% | 85.5% |
| ESM2 8M[1] | LoRA | 99 | 69,2% | 80,7% | 85,4% | 77,0% | 54,7% | 55,5% | 66,0% | 75,9% |
| | | 98 | 69,0% | 77,4% | 83,8% | 81,3% | 59,1% | 56,1% | 66,1% | 76,0% |
| | | 97 | 69,4% | 81,3% | 84,7% | 72,0% | 57,8% | 54,5% | 68,3% | 75,8% |
| ESM2 35M[1] | LoRA | 99 | 69,4% | 77,4% | 87,7% | 75,1% | 59,3% | 58,2% | 68,0% | 78,9% |
| | | 98 | 69,2% | 81,7% | 86,6% | 76,8% | 57,8% | 57,6% | 69,4% | 78,8% |
| | | 97 | 69,1% | 82,5% | 86,6% | 75,8% | 57,2% | 56,5% | 68,5% | 78,7% |
| ESM2 150M[1] | LoRA | 99 | 69,6% | 84,6% | 88,2% | 74,9% | 60,6% | 63,3% | 67,7% | 82,5% |
| | | 98 | 69,6% | 80,7% | 86,8% | 79,9% | 59,0% | 59,6% | 70,6% | 82,5% |
| | | 97 | 69,3% | 85,0% | 87,3% | 79,9% | 60,8% | 59,8% | 69,7% | 82,4% |
| ESM2 650M[1] | LoRA | 99 | 69,1% | 78,6% | 88,0% | 82,8% | 61,7% | 63,9% | 71,1% | 85,6% |
| | | 98 | 69,2% | 84,5% | 89,0% | 85,3% | 64,0% | 63,3% | 72,3% | 85,5% |
| | | 97 | 69,7% | 81,8% | 87,5% | 82,9% | 63,1% | 64,5% | 71,1% | 85,5% |
| ESM2 3B[1] | LoRA | 99 | 69,9% | 85,4% | 89,3% | 84,1% | 71,1% | 68,2% | 71,9% | 86,4% |
| | | 98 | 70,0% | 86,6% | 88,9% | 84,5% | 68,6% | 66,9% | 70,8% | 86,4% |
| | | 97 | 69,9% | 85,0% | 89,8% | 83,6% | 70,1% | 68,4% | 72,0% | 86,5% |
| ProtT5[2] | LoRA | 99 | 66.9% | 77.2% | 87.9% | 80.6% | 72.6% | 66.9% | 70.9% | 84.9% |
| | | 98 | 68.2% | 75.5% | 87.6% | 82.5% | 72.8% | 66.3% | 71.6% | 85.0% |
| | | 97 | 68.8% | 75.0% | 88.1% | 84.0% | 72.4% | 64.5% | 71.3% | 84.9% |
| | | 96 | - | - | - | - | - | 64.5% | 71.7% | 84.9% |
| | | 95 | - | - | - | - | - | 63.3% | 71.1% | 84.8% |
| Ankh base[3] | LoRA | 99 | 69.0% | 83.0% | 86.6% | 77.2% | 62.0% | 63.1% | 68.5% | 84.0% |
| | | 98 | 69.2% | 81.4% | 87.4% | 83.0% | 61.9% | 62.7% | 68.0% | 84.0% |
| | | 97 | 69.3% | 79.6% | 85.5% | 82.6% | 60.1% | 61.8% | 68.8% | 83.9% |
| Ankh large[3] | LoRA | 99 | 69.6% | 84.2% | 87.8% | 80.4% | 57.2% | 61.2% | 69.2% | 85.8% |
| | | 98 | 69.2% | 83.6% | 89.4% | 82.5% | 58.5% | 64.1% | 68.4% | 85.7% |
| | | 97 | 69.8% | 86.1% | 89.2% | 78.2% | 64.0% | 63.1% | 68.3% | 85.6% |

#### 1.2 Aggregated results

**Table S3.** ESM2 - pre-trained embeddings

| | ESM2 8M | ESM2 35M | ESM2 150M | ESM2 650M | ESM2 3B |
|---|---|---|---|---|---|
| GFP | 63.9% ± 0.20 | 65.0% ± 0.16 | 63.9% ± 0.08 | 64.8% ± 0.11 | 65.0% ± 0.14 |
| AAV | 66.9% ± 0.70 | 59.0% ± 3.14 | 65.7% ± 1.63 | 47.1% ± 2.50 | 77.6% ± 0.25 |
| GB1 | 81.8% ± 0.41 | 82.7% ± 0.28 | 83.3% ± 0.95 | 85.7% ± 0.44 | 86.6% ± 0.51 |
| Stability | 75.0% ± 2.02 | 69.1% ± 1.44 | 78.4% ± 1.74 | 66.4% ± 2.67 | 78.9% ± 0.64 |
| Meltome | 56.4% ± 0.46 | 58.6% ± 0.47 | 61.7% ± 0.63 | 65.6% ± 0.56 | 67.7% ± 0.12 |
| Sub. Loc. | 52.0% ± 0.61 | 54.9% ± 0.91 | 59.4% ± 0.92 | 62.9% ± 1.26 | 64.2% ± 0.53 |
| Disorder | 69.7% ± 0.29 | 68.7% ± 0.43 | 71.0% ± 0.48 | 72.0% ± 0.33 | 71.0% ± 0.25 |
| Sec. Str. | 75.3% ± 0.05 | 78.2% ± 0.03 | 82.2% ± 0.06 | 84.7% ± 0.05 | 85.6% ± 0.02 |

**Table S4.** ESM2 - fine-tuning

| | ESM2 8M full model | ESM2 35M full model | ESM2 150M full model | ESM2 650M full model | ESM2 3B LoRA |
|---|---|---|---|---|---|
| GFP | 68.8% ± 0.30 | 68.9% ± 0.24 | 69.1% ± 0.25 | 68.8% ± 0.40 | 69,9% ± 0,08 |
| AAV | 82.7% ± 2.41 | 80.8% ± 3.27 | 84.6% ± 0.37 | 78.3% ± 2.54 | 85,7% ± 0,96 |
| GB1 | 88.3% ± 0.95 | 87.9% ± 0.45 | 87.9% ± 0.19 | 87.9% ± 0.87 | 89,3% ± 0,48 |
| Stability | 76.5% ± 1.96 | 76.9% ± 0.50 | 73.3% ± 1.37 | 74.3% ± 4.89 | 84,1% ± 0,48 |
| Meltome | 58.4% ± 0.40 | 59.8% ± 2.50 | 62.6% ± 1.31 | 66.4% ± 1.47 | 69,9% ± 1,43 |
| Sub. Loc. | 55.9% ± 1.50 | 58.1% ± 1.54 | 61.6% ± 1.16 | 63.8% ± 1.31 | 67,8% ± 0,87 |
| Disorder | 71.5% ± 0.50 | 73.4% ± 0.91 | 74.0% ± 1.10 | 72.0% ± 0.45 | 71,6% ± 0,72 |
| Sec. Str. | 76.1% ± 0.13 | 79.1% ± 0.07 | 82.8% ± 0.10 | 85.5% ± 0.03 | 86,4% ± 0,08 |

**Table S5.** T5 models - pre-trained embeddings

| | ProtT5 | Ankh base | Ankh large |
|---|---|---|---|
| GFP | 61.4% ± 0.42 | 66.0% ± 0.16 | 67.0% ± 0.14 |
| AAV | 68.5% ± 2.45 | 63.7% ± 3.25 | 73.0% ± 1.36 |
| GB1 | 85.2% ± 0.43 | 84.7% ± 0.54 | 85.4% ± 0.73 |
| Stability | 77.8% ± 1.40 | 75.6% ± 2.49 | 69.1% ± 4.20 |
| Meltome | 69.4% ± 0.88 | 57.8% ± 0.76 | 62.0% ± 0.71 |
| Sub. Loc. | 61.3% ± 1.43 | 60.3% ± 0.60 | 60.6% ± 1.28 |
| Disorder | 70.9% ± 0.17 | 71.0% ± 0.25 | 69.8% ± 0.23 |
| Sec. Str. | 84.3% ± 0.05 | 84.4% ± 0.03 | 86.0% ± 0.03 |

**Table S6.** T5 models - fine-tuning

| | ProtT5 LoRA | Ankh base LoRA | Ankh large LoRA |
|---|---|---|---|
| GFP | 68.0% ± 1.10 | 69.2% ± 0.19 | 69.5% ± 0.35 |
| AAV | 75.9% ± 1.35 | 81.3% ± 1.97 | 84.7% ± 1.48 |
| GB1 | 87.9% ± 0.26 | 86.5% ± 1.08 | 88.8% ± 0.95 |
| Stability | 82.4% ± 1.90 | 80.9% ± 3.68 | 80.4% ± 2.44 |
| Meltome | 72.6% ± 0.20 | 61.3% ± 1.19 | 59.9% ± 4.09 |
| Sub. Loc. | 65.1% ± 1.31 | 62.5% ± 0.71 | 62.8% ± 1.64 |
| Disorder | 71.3% ± 0.28 | 68.4% ± 0.50 | 68.7% ± 0.57 |
| Sec. Str. | 84.9% ± 0.07 | 84.0% ± 0.05 | 85.7% ± 0.14 |

#### 2 Detailed evaluation of sub-cellular location and secondary structure prediction

To gain further insights we investigated fine-tuning effect results of both classification tasks (Q10 sub-cellular location prediction and three class per residue secondary structure prediction) on a class level. Results shown here are based on the best ProtT5 predictors for these datasets (Table S2, S1).
For the *Secondary Structure* prediction task (Table S7) the f1-score increased for all three classes. Fine-tuning was not over-fitting to the majority classes, with the largest f1-score gain even seen for the minority class (Strand). While the overall gains were low for this task (refer Fig. 1, Fig. S3, S5) this still was a clear improvement.

**Table S7.** Secondary structure prediction - per class results

| | embedding | | | class share training | fine-tuned | | |
|---|---|---|---|---|---|---|---|
| | precision | recall | f1-score | | precision | recall | f1-score |
| Other | 0.788 | **0.837** | 0.812 | 38.6% | **0.812** | 0.829 | **0.820** |
| Helix | 0.887 | 0.889 | 0.888 | 38.4% | **0.888** | **0.900** | **0.894** |
| Strand | **0.866** | 0.772 | 0.816 | 23.0% | 0.864 | **0.815** | **0.839** |

With more classes and stronger class imbalance the *Sub-cellular location* task was more informative here. The lower performance for underrepresented classes could not be resolved by fine-tuning. Previous findings[4] showed the same tendency. It seems there is not enough training data for these classes in the training data. Still, the significant average improvements (Fig. 1, Fig. S3, S5) together with the balanced improvements seen here (increased or unchanged f1-score for all but the "Plastid" class) are clearly preferable to the embedding based prediction.

**Table S8.** Sub-cellular location prediction - per class results

| | embedding | | | class share training | fine-tuned | | |
|---|---|---|---|---|---|---|---|
| | precision | recall | f1-score | | precision | recall | f1-score |
| Nucleus | 0.632 | **0.848** | 0.724 | 29.0% | **0.788** | 0.828 | **0.808** |
| Cytoplasm | 0.514 | 0.650 | 0.574 | 19.6% | **0.524** | **0.744** | **0.615** |
| Extracellular | 0.896 | 0.750 | 0.817 | 13.9% | **0.901** | **0.793** | **0.844** |
| Mitochondrion | 0.333 | 0.700 | 0.452 | 10.6% | **0.562** | **0.900** | **0.692** |
| Cell membrane | 0.619 | 0.398 | 0.484 | 9.5% | **0.672** | **0.459** | **0.545** |
| Endoplasmic reticulum | 0.467 | 0.412 | 0.437 | 6.3% | **0.548** | **0.500** | **0.523** |
| Plastid | **1.000** | **0.727** | **0.842** | 5.4% | 0.667 | **0.727** | 0.696 |
| Golgi apparatus | **0.333** | 0.154 | 0.211 | 2.5% | **0.333** | **0.231** | **0.273** |
| Lysosome/Vacuole | 0 | 0 | 0 | 2.3% | 0 | 0 | 0 |
| Peroxisome | 0 | 0 | 0 | 1.1% | **0.667** | **0.667** | **0.667** |

These results demonstrated that even in imbalanced datasets, we did not observe over-fitting to majority classes at the expense of minority ones. These findings further reinforce our belief that fine-tuning is a superior approach to using pre-trained embeddings in general.

#### 3 Sub-cellular location prediction results

We provide the results for the *Sub-cellular location* prediction here. Learned light attention (LA) based pooling on pre-trained ProtT5 embeddings had previously[4] shown to be a clear improvement over the use of standard average pooling (ProtT5 FFN). LoRA fine-tuning even surpassed these results without using a complex pooling mechanism like LA.

**Table S9. Q10 sub-cellular location prediction** * Values are taken from previous work[4]. Performance measured as 10-class accuracy (Q10) on the "*set_HARD*" test dataset which estimates the performance for *unkown* proteins that are not sequence-similar to those with experimentally known location.

| Model | set_HARD |
|---|---|
| ProtT5 FFN* | 61.3±1.0 |
| ProtT5 LA* | 65.2±0.6 |
| ProtT5 FFN-LoRA | **66.2±0.6** |

#### 4 GPU requirements for embedding generation

In contrast to fine-tuning (Fig. 5b), embedding computation was much less demanding on GPU memory and therefore can be performed even for datasets with very long sequences (Fig. S1).

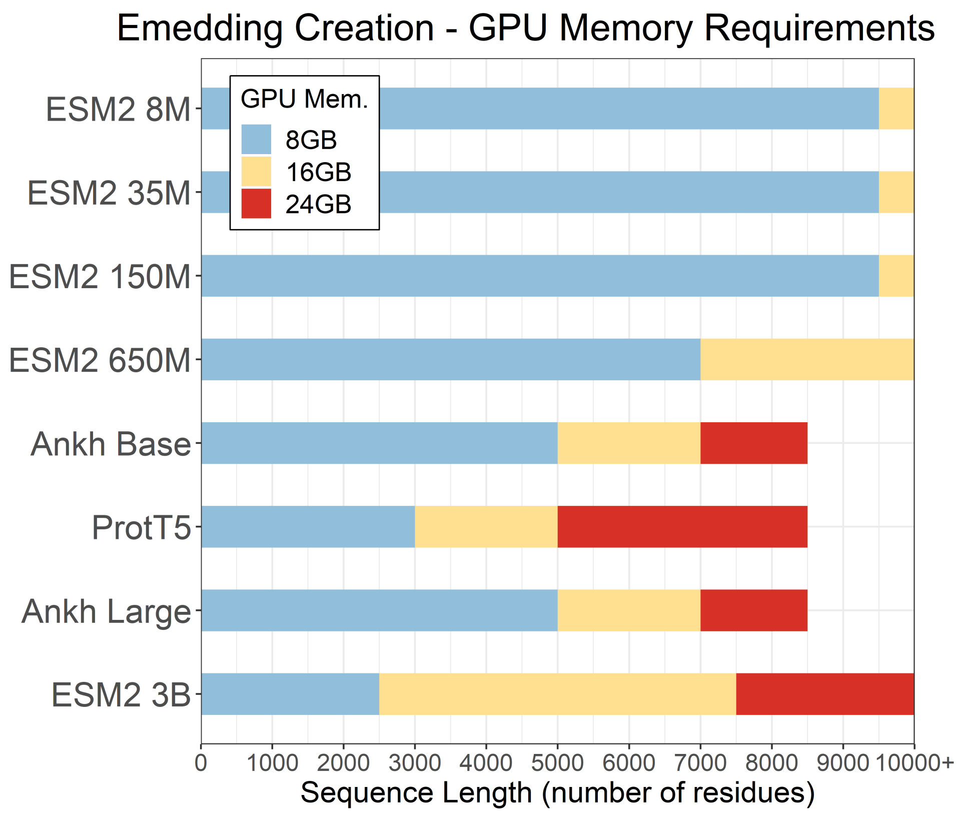


**Figure S1. GPU requirements - embedding creation.** Maximum sequence length before GPU runs out of memory, given a GPU size and model. These values assume the model is loaded in half precision (apart from the Ankh models, which do not support this) and a single embedding is computed at a time.

#### 5 Full model fine-tuning vs LoRA

Analog to the main result heatmap (Fig. 1) we compute the percentage differences between the LoRA fine-tuned and fully fine-tuned models (Eqn.S1) here:

$$\Delta(PT) = performance(PT)_{LoRA} - performance(PT)_{full\_model} \tag{S1}$$

Fig. S2 confirms, that LoRA reaches a similar performance when compared to fine-tuning of the entire model. While there are some exceptions, the difference is not significant for most model / prediction task (PT) combinations. The average improvement across all tasks, shown in the rightmost column, suggests that smaller models should be fully fine-tuned, while for ESM2 650M the use of LoRA fine-tuning is preferable. Contrary to this we decided to still use the full model results for the 650M model (Table S4) to avoid another red (unsuccessful) tile for the *Meltome* task in Fig. 1.
While full fine-tuning of larger models is possible with our available hardware (Fig. 5b), it is not practical due to the resulting significant longer training times (Fig. 5a). Also, during experimentation with ESM2 3B we saw the occurrence of exploding gradients, leading to termination of training runs within the first epoch. While this problem can possibly be mitigated (e.g. deploying lower learning rates or gradient clipping), this decrease in training stability makes full model fine-tuning even less attractive for large models. Due to these findings and the trend seen for the smaller models (Fig. S2), we do not recommend full fine-tuning for PLMs larger than ESM2 650M.

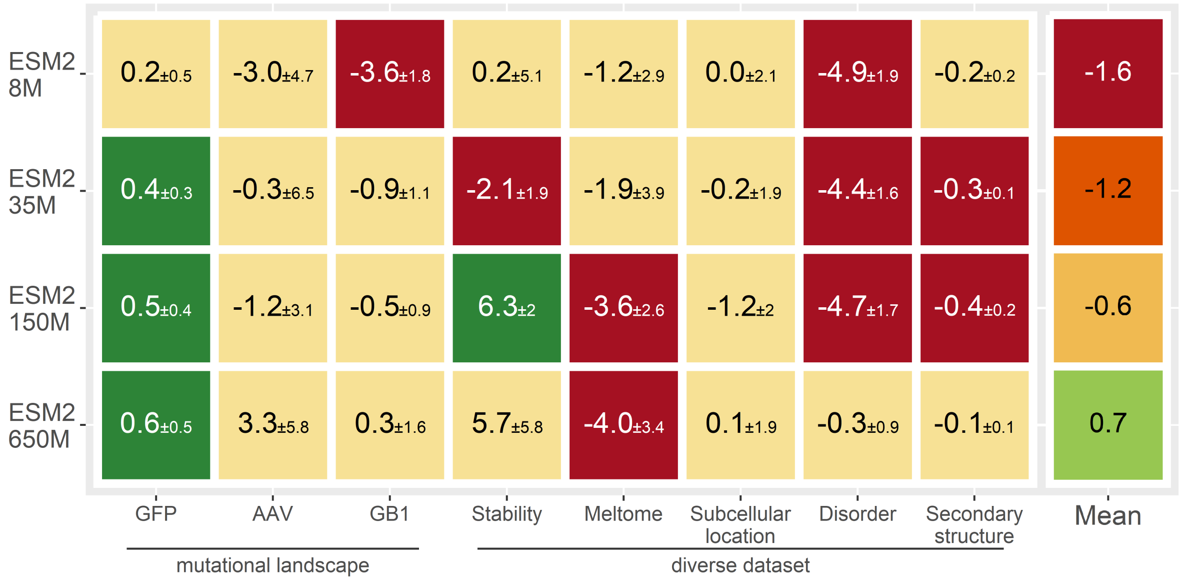


**Figure S2. LoRA compared to fine-tuning full models.** Values represent associated metrics with each dataset (Spearman ranking correlation for *GFP, AAV, GB1, Stability, Meltome* and *Disorder*; accuracy for 10-class, per-protein sub-cellular location and 3-class per-residue secondary structure). Each tile compares LoRA fine-tuning to fine-tuning of the full model for one prediction task. Green tiles mark statistically significant performance increases (exceeding 1.96 standard errors; LoRA over full model fine-tuning), yellow tiles mark statistically insignificant changes (0 lies within the error margins of ±1.96 stderr) and for red tiles LoRA gained significantly less performance. Error estimates (±percentage values) represent the 95% confidence intervals (CI). For individual results see Table S2.

#### 6 Model training - Hyperparameters

We provide the parameters used for model training in Table S10 and S12.

**Table S10.** Training Parameters - pre-trained embeddings

| Dataset | Epochs | Validation per epoch | learning rate | batch size |
|---|---|---|---|---|
| GFP | 240 | 1 | 1e-04 | 8 sequences |
| AAV | 120 | 1 | 1e-04 | 8 sequences |
| GB1 | 240 | 1 | 1e-04 | 8 sequences |
| Stability | 120 | 1 | 1e-04 | 8 sequences |
| Meltome | 120 | 1 | 1e-04 | 8 sequences |
| Sub. Loc. | 120 | 1 | 1e-04 | 8 sequences |
| Disorder | 50 | 1 | 1e-04 | 8 residues |
| Sec. Str. | 10 | 1 | 1e-04 | 8 residues |

**Table S11.** Grid search - Hyperparameter

| | | learning rate | | |
|---|---|---|---|---|
| | | 1e-03 | 3e-04 | 2e-05 |
| | 2 | 64.3% | 63.9% | 65.1% |
| batch | 8 | 65.8% | **66.3%** | 63.3% |
| size | 32 | 65.1% | 64.9% | 47.6% |
| | 128 | 65.7% | 64.7% | 32.2% |

Batch size and learning rate for LoRA fine-tuning were determined using a coarse grained grid search (Table S11), on the sub-cellular location dataset, training the ProtT5 model. We do not expect this to be the most optimal parameter set for all models and datasets but this configuration has shown to be a good starting point leading to stable training behaviour throughout our experiments.

**Table S12.** Training Parameters - fine-tuning

| Model | fine-tuning | Dataset | Epochs | Validation per epoch | learning rate | batch size |
|---|---|---|---|---|---|---|
| ESM2 | full model | GFP | 20 | 5 | 2e-05 | 8 sequences |
| | | AAV | 10 | 5 | 2e-05 | 8 sequences |
| | | GB1 | 20 | 5 | 2e-05 | 8 sequences |
| | | Stability | 10 | 10 | 2e-05 | 8 sequences |
| | | Meltome | 10 | 10 | 2e-05 | 8 sequences |
| | | Sub. Loc. | 10 | 10 | 2e-05 | 8 sequences |
| | | Disorder* | 20 | 10 | 2e-5 / 2e-4 | 1 / 8 sequences |
| | | Sec. Str. | 5 | 20 | 2e-05 | 1 sequence |
| Ankh | LoRA | GFP | 50 | 1 | 3e-04 | 8 sequences |
| | | AAV | 20 | 1 | 3e-04 | 8 sequences |
| | | GB1 | 50 | 1 | 3e-04 | 8 sequences |
| | | Stability | 50 | 1 | 3e-04 | 8 sequences |
| | | Meltome | 10 | 2 | 3e-04 | 8 sequences |
| | | Sub. Loc. | 10 | 10 | 3e-04 | 8 sequences |
| | | Disorder | 20 | 10 | 3e-04 | 1 sequence |
| | | Sec. Str. | 5 | 20 | 3e-04 | 1 sequence |
| ProtT5 ESM2 | LoRA | GFP | 50 | 1 | 3e-04 | 8 sequences |
| | | AAV | 20 | 1 | 3e-04 | 8 sequences |
| | | GB1 | 50 | 1 | 3e-04 | 8 sequences |
| | | Stability | 50 | 1 | 3e-04 | 8 sequences |
| | | Meltome | 20 | 1 | 3e-04 | 8 sequences |
| | | Sub. Loc. | 5 | 10 | 3e-04 | 8 sequences |
| | | Disorder | 20 | 10 | 3e-04 | 1 sequence |
| | | Sec. Str. | 5 | 20 | 3e-04 | 1 sequence |

* for the *Disorder* dataset, fine-tuning full models, ESM2 150M and 650M did not show stable conversion with our default parameters. We had to increase learning rate and batch size to 2e-4 and 8 sequences

#### 7 Model size impact remains dataset dependent

Here we explored the influence of growing model size on prediction performance. Fig. S3 shows this for the eight prediction tasks. While pre-trained embeddings of the largest model (ESM2 3B) were not the best for all individual tasks, we saw an upward slope of the regression line in all cases, which confirms previous findings[5]. Contrary, for fine-tuned models, larger model size only helped performance for the four diverse datasets in the right column, while for the three mutational landscapes size had little to no impact. Interestingly, for the diverse *Disorder* task smaller models were better than larger ones. While the difference between mutational landscapes and the four diverse datasets on the right followed a clear logic, disorder stood out as an exception. We would have expected disorder and secondary structure prediction to behave similarly, as both are per residue tasks, trained on a diverse set of proteins and redundancy reduction is performed in the same way (using mmseqs with the same thresholds). This again shows the need to investigate prediction tasks / datasets individually and common behavior for similar PTs is not guaranteed. In this particular case the key difference might be the dataset size (see SOM Section 10).

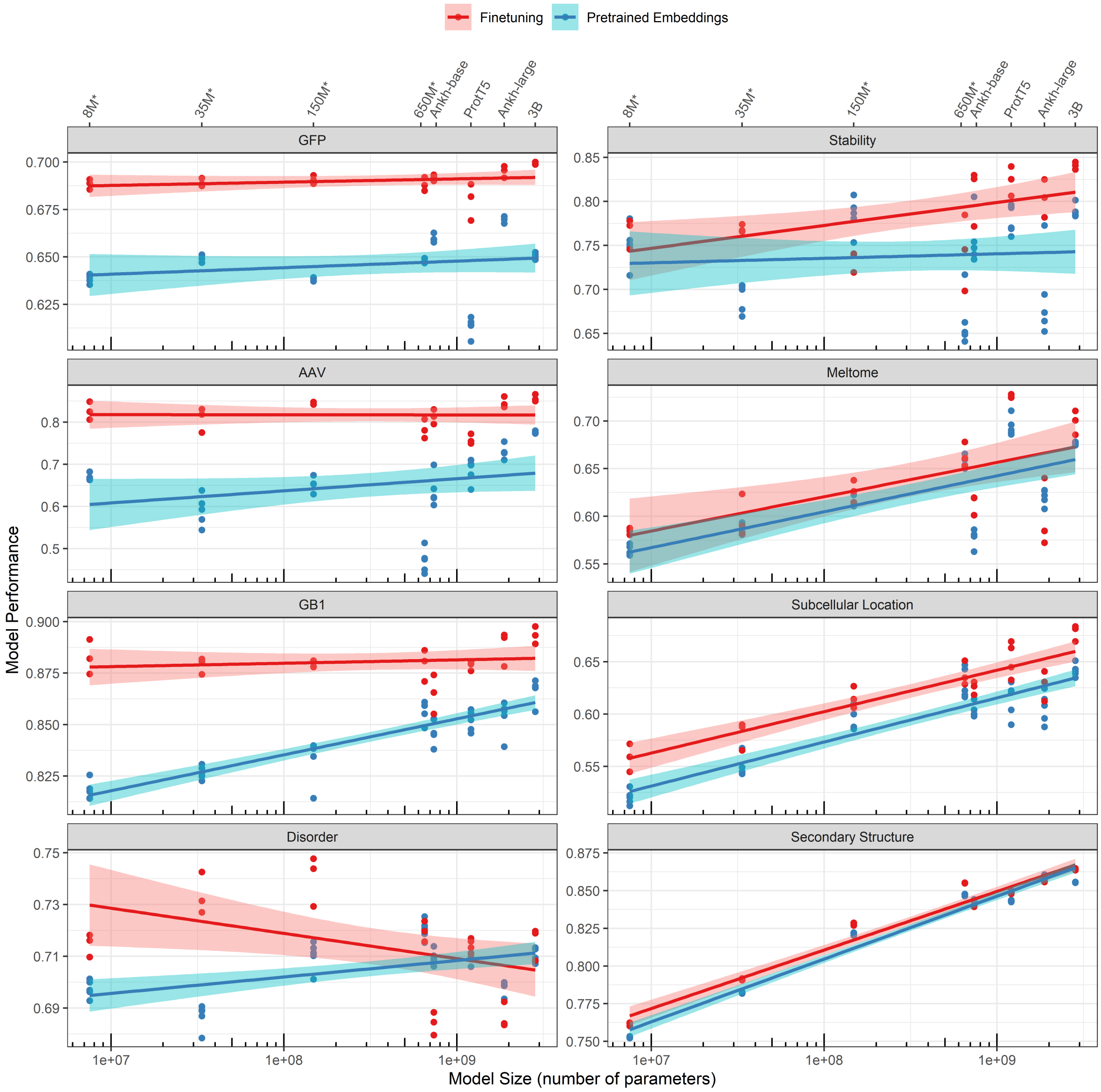


**Figure S3. Model size impact per prediction task.** Reuses the aggregated fine-tuning and pre-trained embedding results from Table S3 - S6, this time mapped to the model size with corresponding models shown at the top. Analog to Fig. 1 and Fig. S5 we report performance of fully fine-tuned models for smaller ESM2 models up to 650M (marked with *). Lighter colored areas are 95% confidence intervals.

#### 8 Initial representation quality important for diverse tasks

Here (Fig.S4) we investigated , whether the quality of initial model representations (pre-trained model embeddings) for individual tasks had influence on the observed fine-tuning gains. For all prediction tasks, but *Meltome*, regression lines showed a downwards slope. This meant that pre-trained models doing well on a specific task will gain less on average compared to other models. This is somewhat intuitive as worse initial performance might allow to pick "lower hanging fruits" making performance increases easier to achieve.

For the mutational landscape datasets (*GFP*, *AAV* and *GB1*), these results are in line with previous findings (Fig. S3), where all fine-tuned models achieved nearly identical performance and differences between models not being significant. This already implied that better initial representations gained less from fine-tuning. Still in two out of three cases the model with best initial representation leads to the best fine-tuned one (Fig. S4 marked as star).

In contrast, for the diverse datasets, the slightly higher gains seen for models with less optimal initial representations do not fully offset this initial gap between representations (Fig. S3). As resulting differences between fine-tuned models are significant here, we suggest to select a model with good initial representation for these tasks.

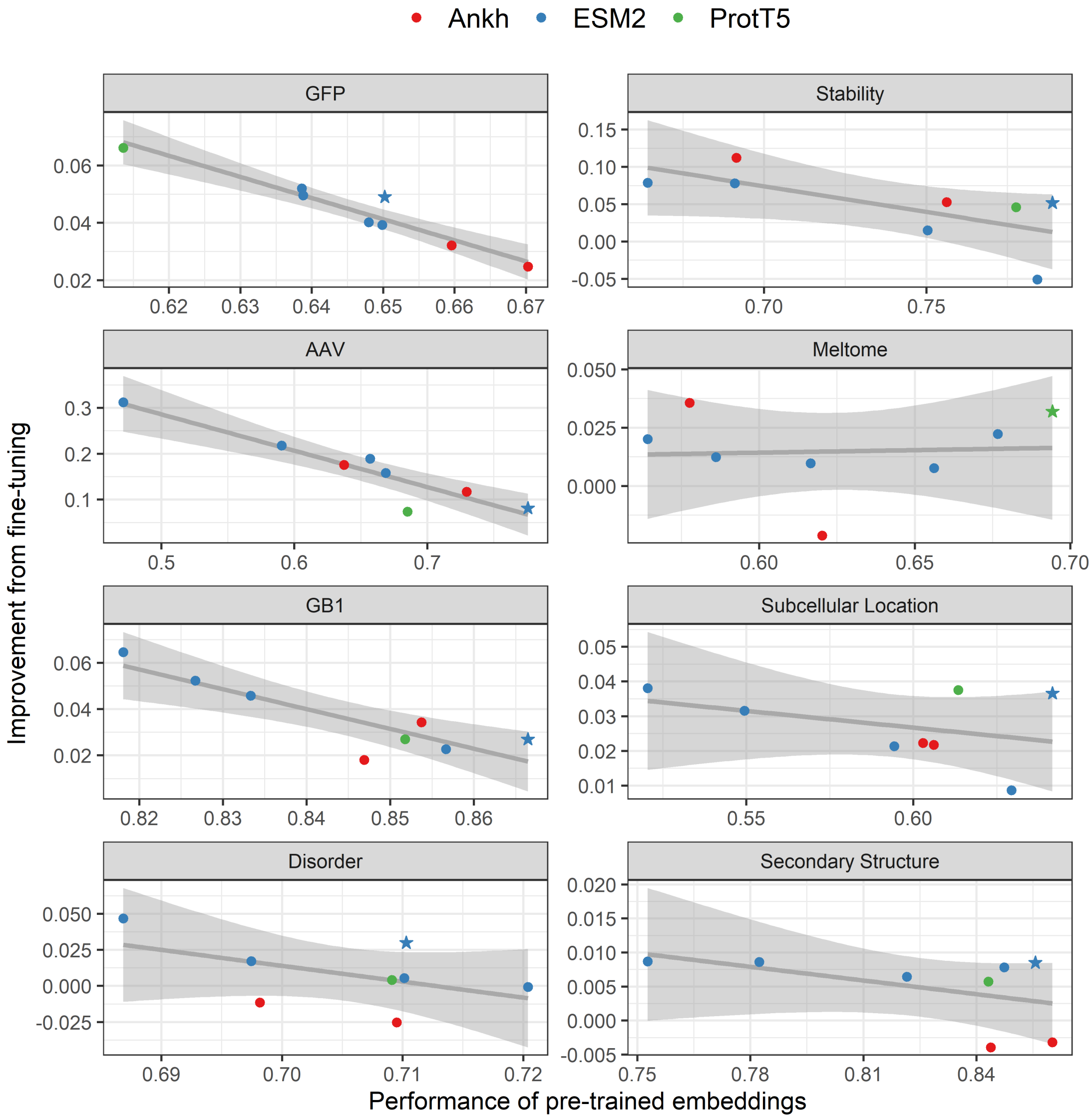


**Figure S4. Impact of initial representation quality.** Reuses the aggregated fine-tuning and pre-trained embedding results from Table S3 - S6. Mapping initial representation quality (performance of pre-trained embeddings) to gains achieved from fine-tuning. The best model (average values) for each task marked as star. Analog to Fig. 1 and Fig. S5 we report performance of fully fine-tuned models for smaller ESM2 models up to 650M. Grey regression line with 95% confidence intervals.

#### 9 Model training behavior

During training we saw three different kind of model behaviours:
First we encountered *noisy test* performance, i.e. the test loss and performance metric are not converging cleanly but stay fluctuating even though training and validation loss flatten out smoothly. This occurred for the *Stability* and *AAV* datasets.
Second, for some tasks we saw *over-fitting*. This occurred for *Meltome, Sub-cellular location* and *Disorder* predictions and is normal behaviour. But for two of those three (*Disorder* and to a lesser extend *Meltome*) the validation loss does not reflected this. Which led to an inability to early stop at the correct epoch.
The third very forgiving behaviour was clean convergence without over-fitting. This happened for the *GFP, GB1* and secondary structure tasks. Here training can be continued beyond initial convergence without any over-fitting.
These behaviours were reflected in the top-level comparison (Fig. 1). To evaluate the impact of these (unwanted) behaviours we analysed the training results a second time. Instead of applying early stopping, we averaged over the 10 highest test performances (which were computed during training parallel to validation performance) for each experiment. This represents a theoretical upper bound of test performance for the predictors (when stopping training at their best performance).

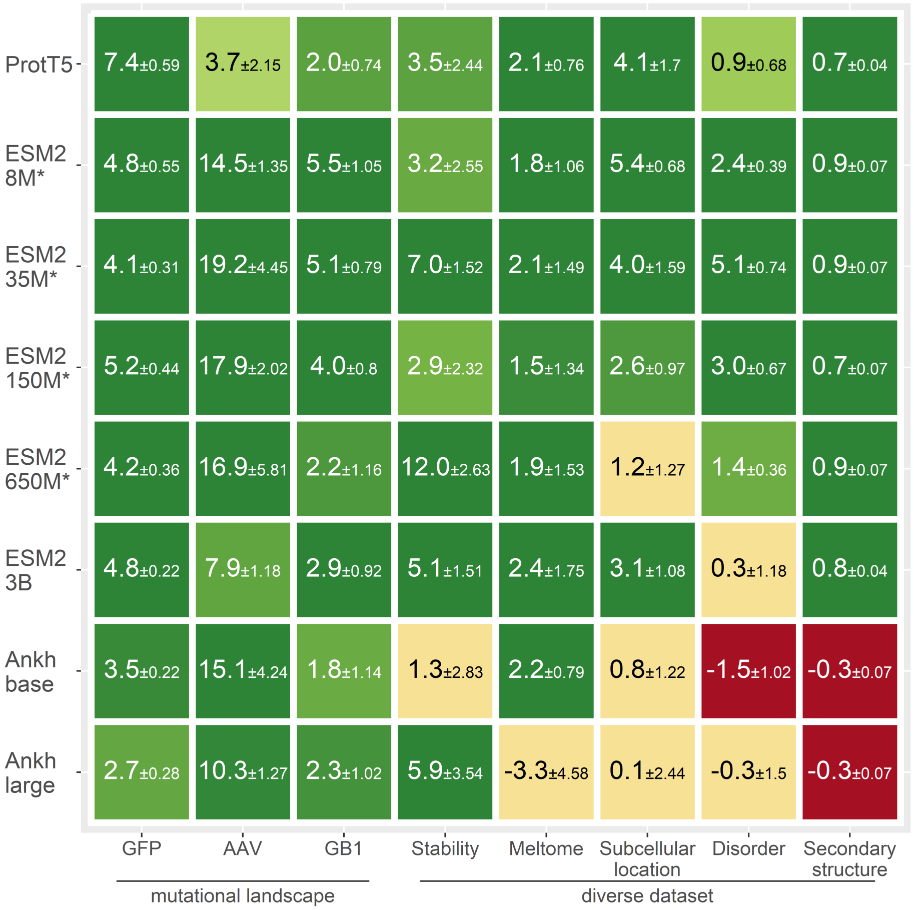


**Figure S5. Upper bound comparison of models and tasks** * marks models which were fully fine-tuned, for the other models we used LoRA (for more details see Fig. S2). Instead of applying early stopping, we averaged over the 10 highest test performance values during training for each individual experiment (per model, task, seed). All further aggregation follows Fig. 1 methodology.

Those upper bound results are shown in Fig. S5 and Tables S13-S16. While the difficulties to fine-tune the Ankh models remained here, supervised fine-tuning looked better for the other models across tasks (compared to Fig. 1). Averaging over the ten best values reduced the random variance from the *noisy test* behaviour, which mainly helped *Stability* (*AAV* improves so much by fine-tuning that the random variance does not matter to much). For the *over-fitting* datasets (*Meltome, Sub-cellular location* and *Disorder*) we also saw improvements. Fine-tuned models train more parameters and were more prone to over-fitting, therefore they gained more compared to embedding based predictors.

##### 9.1 Aggregated results - upper bound

**Table S13.** ESM2 - pre-trained embeddings - upper bound

| | ESM2 8M | ESM2 35M | ESM2 150M | ESM2 650M | ESM2 3B |
|---|---|---|---|---|---|
| GFP | 64.1% ± 0.19 | 65.0% ± 0.17 | 64.0% ± 0.13 | 64.8% ± 0.10 | 64.9% ± 0.11 |
| AAV | 68.6% ± 0.35 | 62.8% ± 1.69 | 67.0% ± 1.40 | 62.7% ± 2.87 | 76.9% ± 0.25 |
| GB1 | 82.9% ± 0.27 | 83.1% ± 0.34 | 84.5% ± 0.42 | 86.3% ± 0.39 | 86.3% ± 0.49 |
| Stability | 76.6% ± 1.45 | 74.0% ± 1.15 | 79.6% ± 1.62 | 70.7% ± 2.27 | 77.6% ± 0.93 |
| Meltome | 57.6% ± 0.18 | 59.0% ± 0.19 | 62.6% ± 0.43 | 66.6% ± 0.39 | 67.2% ± 0.17 |
| Sub. Loc. | 52.8% ± 0.39 | 56.1% ± 0.97 | 60.3% ± 0.64 | 63.9% ± 0.45 | 63.7% ± 0.53 |
| Disorder | 70.0% ± 0.18 | 69.1% ± 0.15 | 71.4% ± 0.12 | 72.3% ± 0.09 | 70.7% ± 0.30 |
| Sec. Str. | 75.2% ± 0.03 | 78.2% ± 0.01 | 82.1% ± 0.02 | 84.6% ± 0.04 | 85.5% ± 0.02 |

**Table S14.** ESM2 - fine-tuning - upper bound

| | ESM2 8M full model | ESM2 35M full model | ESM2 150M full model | ESM2 650M full model | ESM2 3B LoRA |
|---|---|---|---|---|---|
| GFP | 68.9% ± 0.36 | 69.1% ± 0.14 | 69.2% ± 0.31 | 69.0% ± 0.26 | 69,7% ± 0,11 |
| AAV | 83.1% ± 1.00 | 82.0% ± 2.76 | 84.9% ± 0.62 | 79.6% ± 2.94 | 84,8% ± 0,93 |
| GB1 | 88.4% ± 0.78 | 88.2% ± 0.45 | 88.5% ± 0.38 | 88.5% ± 0.77 | 89,2% ± 0,43 |
| Stability | 79.8% ± 1.10 | 81.0% ± 0.37 | 82.5% ± 0.70 | 82.7% ± 0.36 | 82,7% ± 0,58 |
| Meltome | 59.4% ± 0.88 | 61.1% ± 1.30 | 64.1% ± 0.91 | 68.5% ± 1.14 | 69,6% ± 1,58 |
| Sub. Loc. | 58.2% ± 0.29 | 60.1% ± 0.62 | 62.9% ± 0.33 | 65.1% ± 0.82 | 66,8% ± 0,55 |
| Disorder | 72.4% ± 0.21 | 74.2% ± 0.59 | 74.4% ± 0.55 | 73.7% ± 0.27 | 71,0% ± 0,88 |
| Sec. Str. | 76.1% ± 0.04 | 79.1% ± 0.06 | 82.8% ± 0.05 | 85.5% ± 0.03 | 86,3% ± 0,02 |

**Table S15.** T5 models - pre-trained embeddings - upper bound

| | ProtT5 | Ankh base | Ankh large |
|---|---|---|---|
| GFP | 61.6% ± 0.29 | 66.1% ± 0.14 | 67.1% ± 0.15 |
| AAV | 72.4% ± 0.61 | 66.1% ± 1.92 | 74.4% ± 0.57 |
| GB1 | 86.0% ± 0.34 | 85.1% ± 0.36 | 86.5% ± 0.23 |
| Stability | 79.8% ± 0.74 | 80.0% ± 0.89 | 75.8% ± 2.20 |
| Meltome | 70.3% ± 0.34 | 58.4% ± 0.30 | 62.6% ± 0.58 |
| Sub. Loc. | 61.9% ± 1.25 | 61.1% ± 0.64 | 61.4% ± 0.73 |
| Disorder | 71.3% ± 0.16 | 71.0% ± 0.20 | 70.1% ± 0.09 |
| Sec. Str. | 84.2% ± 0.01 | 84.3% ± 0.02 | 86.0% ± 0.02 |

**Table S16.** T5 models - fine-tuning - upper bound

| | ProtT5 LoRA | Ankh base LoRA | Ankh large LoRA |
|---|---|---|---|
| GFP | 69.0% ± 0.30 | 69.6% ± 0.08 | 69.8% ± 0.13 |
| AAV | 76.1% ± 1.54 | 81.2% ± 2.32 | 84.7% ± 0.70 |
| GB1 | 88.0% ± 0.40 | 86.9% ± 0.78 | 88.8% ± 0.79 |
| Stability | 83.3% ± 1.70 | 81.3% ± 1.94 | 81.7% ± 1.34 |
| Meltome | 72.4% ± 0.42 | 60.6% ± 0.49 | 59.3% ± 4.00 |
| Sub. Loc. | 66.0% ± 0.45 | 61.9% ± 0.58 | 61.5% ± 1.71 |
| Disorder | 72.2% ± 0.52 | 69.5% ± 0.82 | 69.8% ± 1.41 |
| Sec. Str. | 84.9% ± 0.03 | 84.0% ± 0.05 | 85.7% ± 0.05 |

#### 10 Data saturation

To gain a better understanding on why over-fitting (SOM Section 9) occurred, we investigated how fine-tuned model performance changed when trained only on a random subset of the original datasets. For these experiments we started with the entire dataset and gradually halved the data again and again until reaching 3.125% of the original size. To keep training times reasonable we chose the ESM2 150M model and again reran each experiment three times. We excluded *AAV* and *Stability* here as both did not even show stable training behavior for their full dataset.

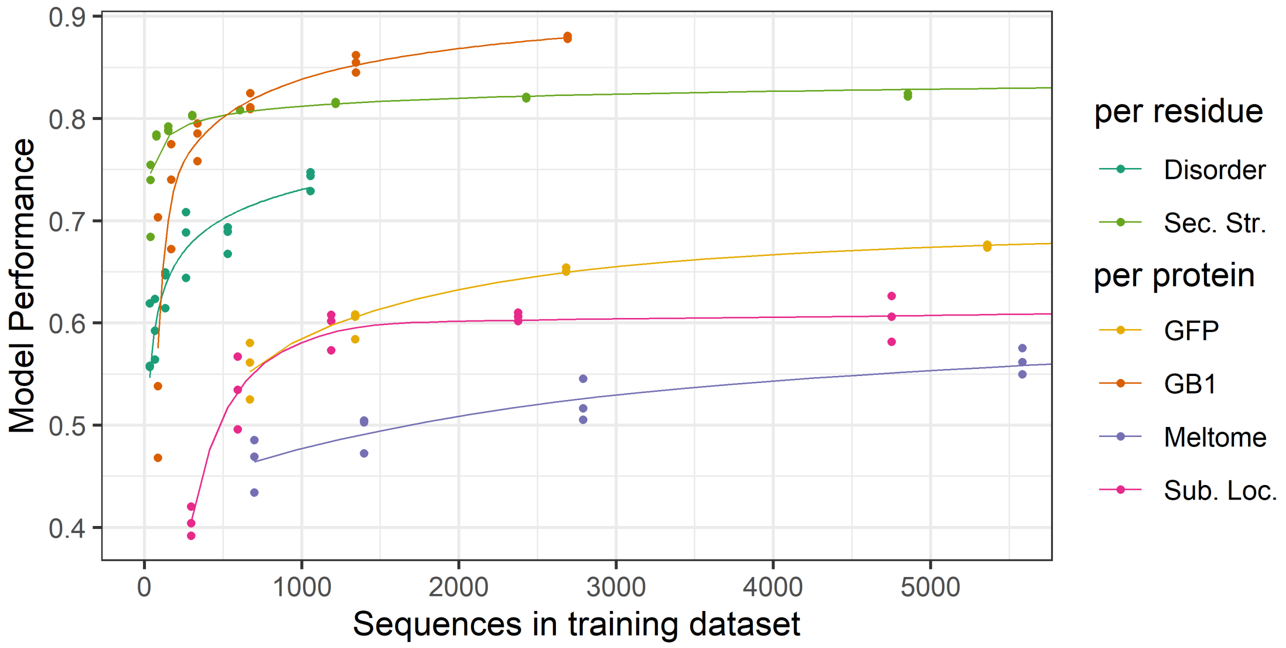


**Figure S6. Downsampled datasets** This shows fine-tuned (full model) ESM2 150M results, trained on randomly down-sampled datasets, each experiment rerun three times. Values represent associated metrics with each dataset (Spearman ranking correlation for *GFP, GB1, Meltome* and *Disorder*; accuracy for 10-class, per-protein sub-cellular location and 3-class per-residue secondary structure). The complete results, including data-points above the x-axis threshold shown here, can be found in the source data

Fig. S6 shows the results. For the *Sec.Str.* and the *GFP* datasets which showed no over-fitting tendency before, we saw an over saturation, meaning reducing the dataset to half or even on fourth (in case of *Sec.Str.*) did not reduce predictor performance. For the third dataset not prone to over-fitting, *GB1*, a further increase in dataset size is not possible by design. All possible single, double and triple mutants are already in the training set and the predictor tries to generalize to test variants where all four selected positions are mutated. The upward slope showed that the predictors benefited from every added lower order variant, confirming the strong epistatic effects between those four positions[6].

The *Disorder* dataset would likely benefit from more data. It is also the smallest dataset shown here, which explains the over-fitting tendency as well as the comparatively worse performance of larger models (as those are even more prone to over-fit) on this dataset (see Fig. S3).

*Sub-cellular* location prediction was not gaining a lot from more data. This was driven by the strong class imbalance (Table S8) and the lack of data for minority classes during training. There was enough data for majority classes (therefore down sampling did not hurt) but too little data on minority classes (that's why overall accuracy is relatively low). This imbalance also caused the over-fitting seen for this dataset. When training was continued for too long the models started to over-fit to majority classes (data not shown).

While *Meltome* was the largest dataset, we still found a performance increase up to the full data set size. As shown before (Fig. S3), *Meltome* favored larger models, which likely meant the sequence-function (thermo-stability) relationship is too complex for smaller models to capture. We were not able to track down the cause of over-fitting here, but found that over-fitting weakened with model size and the two best performing models (ESM2 3B and ProtT5) were hardly affected.

#### 11 ESM2 fine-tuning with random initialization

To determine how much of the performance gains achieved by fine-tuning (over the use of frozen embeddings) can be attributed to the model architecture itself and how much to the model's pre-training, we fine-tuned randomly initialized models.
For training we randomly initialized all model parameters (embedding weights, model weights and the prediction head) and reused the training scripts from our intra-model comparison. Due to the random starting point model training took much longer to converge. We ran each training for 250 epochs and only validated once per epoch.

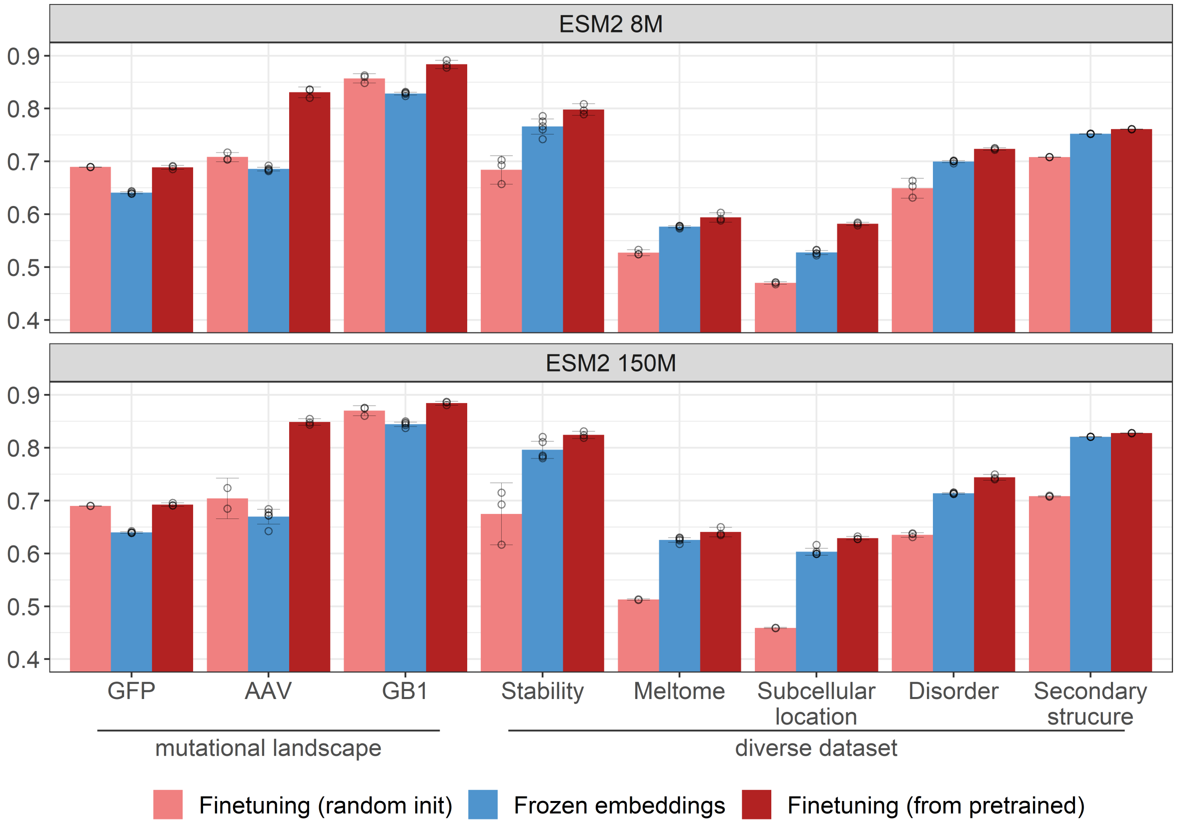


**Figure S7. Model fine-tuning from random initialization.** Values represent associated metrics with each dataset (Spearman ranking correlation for *GFP, AAV, GB1, Stability, Meltome* and *Disorder*; accuracy for 10-class, per-protein sub-cellular location and 3-class per-residue secondary structure), error bars mark the 95% confidence intervals (CI), calculated from multiple reruns of the same model training. Circles represent individual training runs.

We find (Fig. S7, Table S17) similar results for both tested models (ESM2 8M and 150M). For all five diverse datasets we see that models fine-tuned from fully randomly initiated weights (light red) do not reach the performance of predictors using pre-trained embeddings (blue). We concluded that for these datasets pre-training provides a major contribution.
For the mutational landscapes frozen embeddings performed worse than models fine-tuned from random initialization. In case of *GFP* we even found no difference between fine-tuning from pre-trained models compared to starting from random weights. This explains previous findings[7] about the lack of discriminating power of the TAPE[8] *GFP* datasplit. While fine-tuning from pre-trained models still preformed better for the other two landscapes and is also recommended due to much faster training convergence (at least 10 times faster, depending on the dataset), random models did surprisingly well here.
We draw from these results that transfer learning from the unsupervised pre-training is less effective for local landscapes and a large portion of model performance stems from the transformer architecture itself.

**Table S17.** Detailed results - Model fine-tuning from random initialization

| | Fine-tuning (random init) | Frozen embeddings | Fine-tuning (from pre-trained) |
|---|---|---|---|
| **ESM2 8M** | | | |
| GFP | 68.9% ± 0.02 | 64.1% ± 0.2 | 68.9% ± 0.4 |
| AAV | 70.8% ± 0.9 | 68.6% ± 0.4 | 83.1% ± 1.0 |
| GB1 | 85.7% ± 0.8 | 82.9% ± 0.3 | 88.4% ± 0.8 |
| Stability | 68.4% ± 3 | 76.6% ± 1.4 | 79.8% ± 1.0 |
| Meltome | 52.7% ± 0.5 | 57.6% ± 0.2 | 59.4% ± 0.9 |
| Sub. Loc. | 47% ± 0.2 | 52.8% ± 0.4 | 58.2% ± 0.3 |
| Disorder | 64.9% ± 2 | 70.0% ± 0.2 | 72.4% ± 0.2 |
| Sec. Str. | 70.8% ± 0.02 | 75.2% ± 0.03 | 76.1% ± 0.04 |
| **ESM2 150M** | | | |
| GFP | 69% ± 0.03 | 64.0% ± 0.1 | 69.2% ± 0.3 |
| AAV | 70.4% ± 4 | 67.0% ± 1.4 | 84.9% ± 0.6 |
| GB1 | 87% ± 0.9 | 84.5% ± 0.4 | 88.5% ± 0.4 |
| Stability | 67.5% ± 6 | 79.6% ± 1.6 | 82.5% ± 0.7 |
| Meltome | 51.3% ± 0.1 | 62.6% ± 0.4 | 64.1% ± 0.9 |
| Sub. Loc. | 45.9% ± 0.1 | 60.3% ± 0.6 | 62.9% ± 0.3 |
| Disorder | 63.5% ± 0.4 | 71.4% ± 0.1 | 74.4% ± 0.6 |
| Sec. Str. | 70.8% ± 0.1 | 82.1% ± 0.02 | 82.8% ± 0.05 |
